# A Transcriptomic Atlas of Macaúba Palm Reveals Organ-Specific Gene Expression and Stress-Related Pathways

**DOI:** 10.1101/2025.06.27.661789

**Authors:** Lucas Miguel de Carvalho, Bárbara Regina Bazzo, Camila Carlos-Shanley, Carlos Augusto Colombo, Gonçalo Amarante Guimarães Pereira, Marcelo Falsarella Carazzolle

## Abstract

The macaúba palm (*Acrocomia aculeata*) is an emerging oilseed species with promising applications in biodiesel production, as well as in food and cosmetic industries. Native to the Neotropical, it is incipiently domesticated and distributed across diverse environments and edaphoclimatic conditions. However, genomic studies of macaúba are limited due to the scarcity of publicly available sequence data, as it is considered a non-model plant. In this study, we present an exploratory analysis of a transcriptome dataset comprising seven organs (roots, bulbs, male and female flowers, leaves, leaf sheath, and fruits). A total of 22,703 transcripts were assembled into a single reference dataset. Of these, 9,729 transcripts (42.85%) were annotated using KEGG orthology. Gene expression profiling revealed 306, 32, 41, 159, 158 and 916 organ-specific transcripts in leaves, leaf sheaves, bulbs, flowers, fruit and root, respectively. Comparative analysis with African oil palm (*Elaeis guineensis*) and date palm (*Phoenix dactylifera*) revealed 55 gene families exclusive to macaúba palm. In addition, 221 transcripts related to drought stress were identified, grouped into 112 families. Root libraries revealed 7,091 fungal transcripts - approximately 3.9% of all reads – mainly derived from arbuscular mycorrhizal fungi (AMF) Rhizophagus spp. These findings highlight the central role of signal transduction pathways in response to environmental stresses in macaúba palm. The transcriptome dataset generated in this study provides a valuable genomic resource for future genotype-phenotype investigations in macaúba palm. Furthermore, the presence of AMF-associated transcripts suggests a potentially important role for these symbiotic fungi in macaúba palm growth and development.

## 1. Introduction

The oleaginous macaúba palm (*Acrocomia aculeata* (Jacq.) Lodd. Ex Mart., 2n = 30), a native species of Neotropical America belonging to the Arecaceae family, is gaining recognition as a promising oil-feedstock crop (Azevedo Filho et al., 2012; Colombo et al., 2018). In contrast to the oil palm, which is linked to significant sustainability issues such as deforestation, loss of biodiversity, and social conflicts due to its high-water demand (Fitzherbert et al., 2008; Moreira et al., 2020; Rival and Levang, 2014; Sheil et al., 2009), macaúba palm represents an environmentally and socially co-beneficial biofuel source. Additionally, it offers an alternative food source for the human diet in South America (Toledo e Silva et al., 2022).

Macaúba palm grows naturally in large populations, and its presence has been documented from north Florida, Mexico and West Indies to south Paraguay and north Argentina. Its oldest fossil sites are dated to 11,200 years BC in Santarém, Pará - Brazil (as *Acrocomia sp*.), followed by Panama (8,040 B.P.), and Mexico (6,750 B.P.) (Morcote-Ríos and Bernal, 2001). In Brazil, it is the most widely distributed palm, predominantly found in the Cerrado (Brazilian savanna) but also inhabiting arid and semi-arid regions. This extensive distribution reflects its adaptability to diverse soil and climatic conditions, suggesting morphological and molecular adaptations to different environments, which likely contribute to the high genetic variability observed within the species (Díaz et al., 2021).

The macaúba palm exhibits significant morphological and physiological adaptations to a variety of environmental conditions, including irregular precipitation. These adaptations include a specialized root system, photosynthetic adjustments, rapid recovery from stress, and the accumulation of reserves to support seedling growth (Colombo et al., 2018; Marques et al., 2017; Pires et al., 2013; Rosa et al., 2019). The root system plays a crucial role in plant growth and development under both abiotic and biotic stress, directly interacting with the soil and exhibiting developmental plasticity. This adaptability enables plants to respond effectively to diverse environmental conditions and habitats, particularly under water scarcity. Root evolution is an important factor in the advent of land plants and contributed to CO_2_ reduction in Silurian and Devonian (Kenrick and Crane, 1997; Shekhar et al., 2019). In macaúba palm, studies have shown that roots are initially concentrated in the tuberous region in young plants and become uniformly distributed in adult plants, with the effective root depth increasing with age and corresponding to the crown projection area (Moreira et al., 2019, 2018).

Genomic and transcriptomic studies could provide valuable insights into the molecular mechanisms underlying the macaúba palm’s adaptability to diverse soil and climatic conditions. Additionally, de novo transcriptome analysis is particularly useful for plants without a reference genome, and it has been extensively applied to study various palm species. For example, transcriptome data from African oil palm have been used to elucidate host-pathogen interactions (Ho et al., 2016), waterlogging response in two cultivars (Nuanlaong et al., 2020), and lipid biosynthesis mechanisms in oil-storing tissues during developmental stages (Dussert et al., 2013). (Zhang et al., 2012) produced a comprehensive transcriptomic atlas of date palm (*Phoenix dactylifera*, L.) in different organs (roots, branches, male and female flowers, leaves and young and mature fruits) and found organ-specific gene families.

Despite the growing commercial interest in macaúba palm, genomic studies are limited and have primarily focused on population genetics (Bazzo et al., 2018; Lopes et al., 2018). In this study, we present a comprehensive transcriptomic atlas of the macaúba palm, aiming to establish a foundation for the functional genomics of this species to facilitate its breeding and domestication. RNA-Seq transcriptome analysis revealed 22,703 transcripts in macaúba palm (*Acrocomia aculeata*) across seven organs: leaves, leaf sheaths, roots, bulbs, fruit pulp, male flowers, and female flowers. We analyzed organ-specific transcripts, abundance levels, and expression patterns of genes involved in key biological processes. Additionally, we examined gene family expansion compared to other palm genomes and explored genes associated with drought tolerance.

## 2. Material and Methods

### 2.1 Plant Material and RNA Isolation

The plant material used in this study is the same as that used by Bazzo et al., 2018, which are deposited at NCBI in the BioProject PRJNA489676. In their work, Bazzo et al., (2018) analyzed this dataset to develop expressed sequence tag (EST) markers for assessing genetic diversity in macaúba. In the present work, we focus on the functional analysis of this dataset.

For transcriptome sequencing, we used leaves (pinna), leaf sheaves (petioles), roots, bulbs, pulp fruit, and both male and female flowers from macaúba palm (*Acrocomia aculeata*).

Vegetative samples of macaúba palm were collected from eight-month-old seedlings of native plants originating from cities of Dourado, Amparo, and Pedreira (São Paulo State/Brazil). One seedling per location was used, totaling three biological replicates for each vegetative organ (leaves, leaf sheaths/petioles, roots, and bulb). This location diversity was intended to capture a broader range of gene expression patterns representative of different environmental conditions. At eight months of age, the seedlings exhibit well-differentiated vegetative tissues, including leaves, petioles, and roots, as well-developed bulb tissue, allowing comprehensive transcriptomic analysis across organ types. The plants were acclimated for one month in a greenhouse with artificial irrigation conditions.

Female and male flowers were obtained from three adult plants grown at the Santa Elisa Experimental Unit - IAC (Agronomic Institute of São Paulo, Campinas, São Paulo State, Brazil), each plant representing one biological sample.

Pulp fruit samples were collected in natural populations located in Santo Antônio de Posse and Amparo (Sao Paulo State, Brazil), and Ibituruna (Minas Gerais State, Brazil), which each plant from each location representing one biological sample.

Total RNA was extracted using the lithium chloride method (Le Provost et al., 2007; Zeng and Yang, 2002). As fruit tissues present a high content of polyphenols and oil, which interfere in most nucleic acid extraction methods, RNA isolation was performed using the perchlorate protocol described by Rezaian and Krake (1987). RNA quality and concentration were assessed using a 2100 Bioanalyzer (Agilent Technologies, Santa Clara, CA) and NanoDrop 2000 spectrophotometer (Thermo Fisher Scientific Inc.).

### 2.2 De novo transcriptome sequencing, assembly, quantification and annotation

All RNA-Seq reads generated from all different organs were deposited in the NCBI Sequence Read Archive (SRA), under BioProject PRJNA489676. mRNA libraries were prepared using the TruSeq Stranded mRNA Library Preparation Kit, and sequencing was carried out on the Illumina HiSeq™ 2500 platform with 100-bp paired-end reads (Illumina, San Diego, CA, USA). Quality control of raw reads was performed using FasTQC software (Andrews, 2010). Ribosomal RNA reads were discarded using SortMeRNA (Kopylova et al., 2012), and remaining reads were assembled using *de novo* transcriptome assembler Trinity version 2.0.6 (Grabherr et al., 2011).

Transcript quantification (TPM - Transcripts per Million) was estimated using Kallisto v 0.44.0 (Bray et al., 2016), and the expression was evaluated at both the transcript and gene level. For gene-level quantification, expression values corresponded to the sum of all transcripts originating from the same locus (sum of TPMs). Open reading frames (ORFs) were identified using the TransDecoder program v4.1.0 (https://github.com/TransDecoder/TransDecoder) configured to retain only the largest ORF with at least 67 codons (201 nt). Since the presence of multiple transcripts per locus (due to alternative splicing events), the final transcriptome was refined and obtained by considering only one transcript representative of each locus (longest ORF), with a gene-level TPM ≥ 1 in at least one of the analyzed organs.

Functional annotation was conducted using the PANNZER software v.1.0 (Koskinen et al., 2015) and the NCBI/CDD database (Marchler-Bauer et al., 2011), considering an expect value threshold of 1e-10, extracting only the top hit for each sequence. Non-plant transcripts were identified and removed using NCBI Foreign Contamination Screen (FCS) tool (Astashyn et al., 2024). The ExpressionAnalyst platform was used to perform principal component analysis (PCA), pairwise comparisons and over-representation analysis (Ewald et al., 2023).

### 2.3 Organ-specificity analysis

To identify the set of organ-specific genes, the SPM (specificity measure) metric was applied for each gene using the software tspex (Camargo et al., 2020), only SPM >= 0.95 were considered (Xiao et al., 2010).

### 2.4 Orthologous analysis and comparative genomes

Protein sequences of *Elaeis guineensis, Musa acuminata, Phoenix dactylifera and Oryza sativa* (used as an outgroup) were obtained from the Phytozome V12 repository (https://genome.jgi.doe.gov/). The sequences, along with the proteins from the macaúba transcriptome (longest ORF translated) were used to identify ortholog groups using the program OrthoFinder v1.1.8 (Emms and Kelly, 2015).

A total of 1,994 single-copy orthologs were aligned with MAFFT v.7.310 (Katoh et al., 2002) with maximum number of iterations set to 1000. Maximum likelihood phylogenetic analysis was performed using the RAxML software v.8.2.11 (Stamatakis, 2014). Bootstrap values were generated with 1.000 replicates. Gene family expansion analysis in macauba was performed with BadiRate v.1.35 (Librado et al., 2012) applying an FDR threshold of 0.05. To identify high expressed transcripts associated to hydric stress, we compared our orthogroups data with Drought DB – Drought Stress Gene Database (Alter et al., 2015). We performed a BLASTp to compare the Drought DB database with our *de novo* transcriptome (translated ORFs), using an e-value cutoff of 1e-05.

The transcriptome Atlas pipeline from macaúba RNA-Seq reads described in this study are summarized in Figure 1.

**Figure 1.**
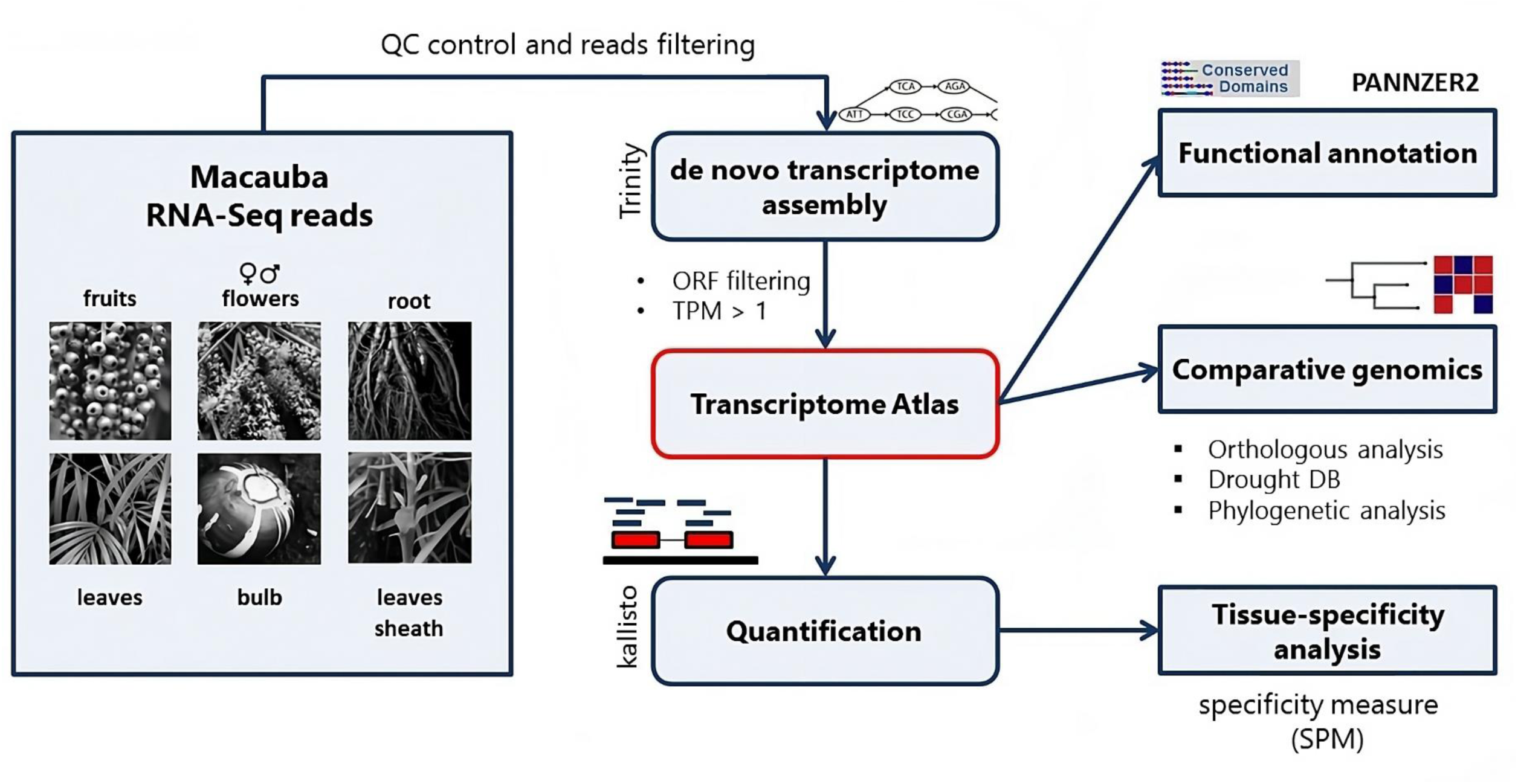
Transcriptome Atlas analysis pipeline from macaúba organs RNA-Seq reads.

## 3. Results

### 3.1 Transcriptome assembly and annotation

A total of 567 million 100-bp paired-end reads were generated from sequencing the transcriptomes of macaúba organs - leaves, leaf sheaths, bulbs, roots, female flowers, male flowers, and fruits - with an average of 27 million reads per organ. After quality control and the removal of rRNA reads, 522 million high-quality reads (92.08%) were retained for de novo assembly. This process yielded 34,293 transcripts, with an N50 length of 1,483 bp; the longest transcript measuring 8,698 bp, and the shortest at 297 bp. Among these, 7,091 transcripts were assigned as fungi, and 4,499 were removed using a cutoff of minimum TPM of 1 in at least one organ.

A total of 22,703 macaúba palm transcripts were used in our analyses (Supplementary File 1) of which 2,803 transcripts (12,34%) were successfully annotated using UniProtKB/Swiss-Prot, 18,020 transcripts (79.37%) were annotated with CDD, 4,129 transcripts (18.18%) were associated with at least one Gene Ontology (GO) term from the three main GO categories, and 9,729 transcripts (42.85%) were annotated using KEGG orthology (KO).

Principal component analysis (PCA) of the KO functions revealed that PC1 effectively separated the transcriptomes of all organs except the bulb, which overlapped with the flowers and roots. PC2 primarily distinguished the transcriptome of ripe fruits (Figure 2).

**Figure 2.**
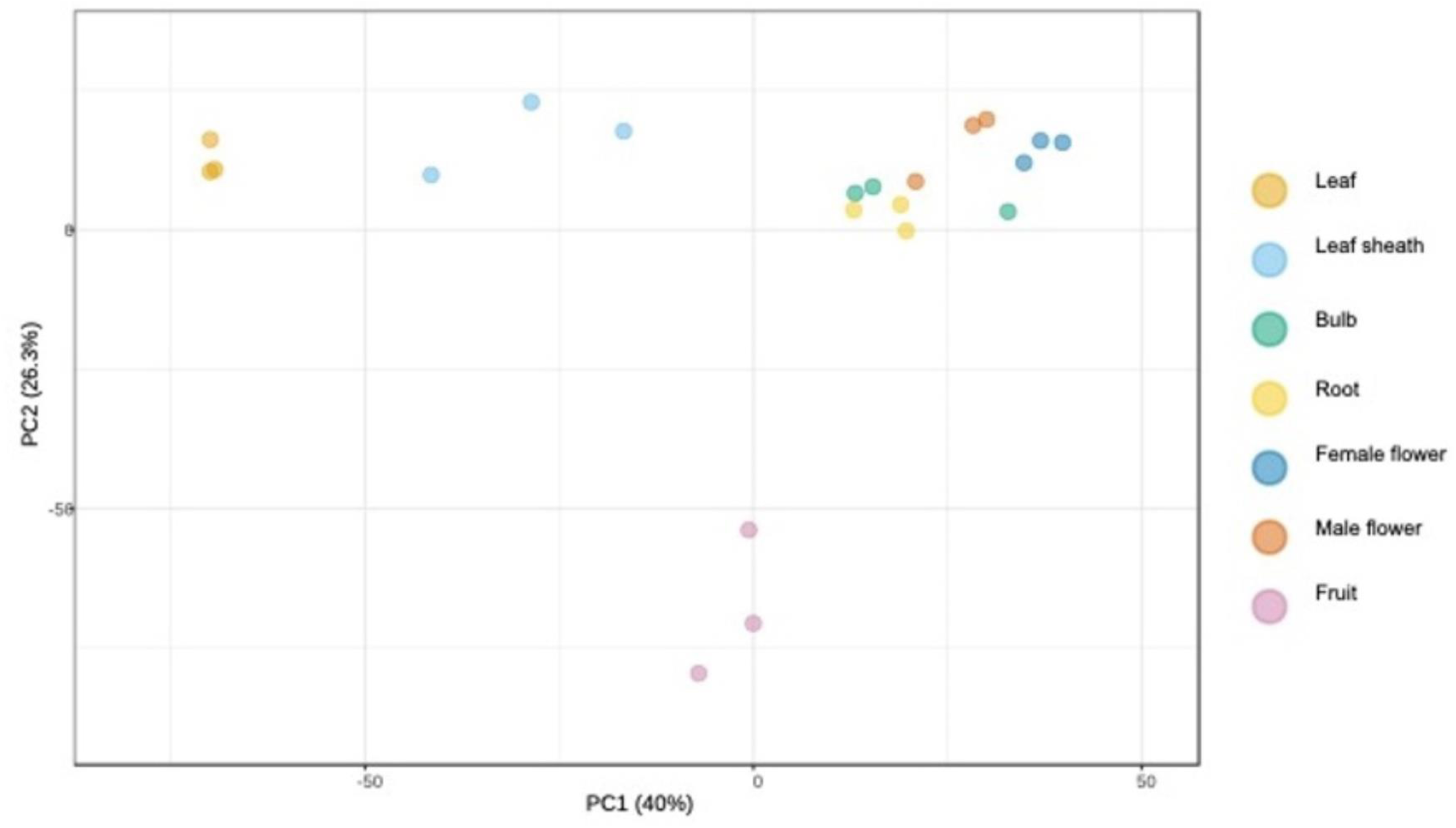
Principal component analysis (PCA) based on the Transcripts-per-Million (TPM) values of KEGG Orthogy (KO) functions.

Pairwise comparisons revealed organ-specific upregulation patterns. For example, genes involved in the Peroxisome pathway were more highly expressed in the leaf sheath compared to the leaf, whereas genes involved with carbon fixation and glyoxylate/dicarboxylate metabolism were more expressed in the leaf (Supplementary Figure 1). The comparison between root and bulb organ revealed that genes related to fatty acid and Nitrogen metabolism were more upregulated in the root (Supplementary Figure 2). In female versus male flowers, genes involved in the cell cycle and DNA replication pathways showed higher expression in the female flowers (Supplementary Figure 3). Additionally, a comparison between female flowers and fruits demonstrated increased expression of genes associated with fatty acid and Starch and sucrose metabolism in the fruits (Supplementary Figure 4).

### 3.2 Top expressed transcripts

The ten most highly expressed transcripts in each were ranked organ based on TPM values (Figure 3). As expected, the top transcripts in leaves and leaf sheaths were related to photosynthesis. Notably, the most highly expressed transcript in both female and male flower was a hypothetical protein (TRINITY_DN35918_c5_g2), with a transcript size of 961 bp encoding a 313 amino acid protein, and median TPM value of 21,747.01 and 37,996.77 for female and male flower, respectively. In the bulb, the most expressed transcript was tubulin alpha (TRINITY_DN35302_c5_g1) with a median TPM of 9,967.61. Overall, bulb and root exhibited similar expression profiles, suggesting a potential correlation between these underground organs, which are subject to the same demands for storage, nutrient transport, or stress response. Additionally, a highly expressed transcript (TRINITY_DN22886_c0_g1) was identified in root, bulb and leaf sheath, encoding a dormancy-associated protein, an auxin-repressed involved in the regulation of plant growth and development.

**Figure 3.**
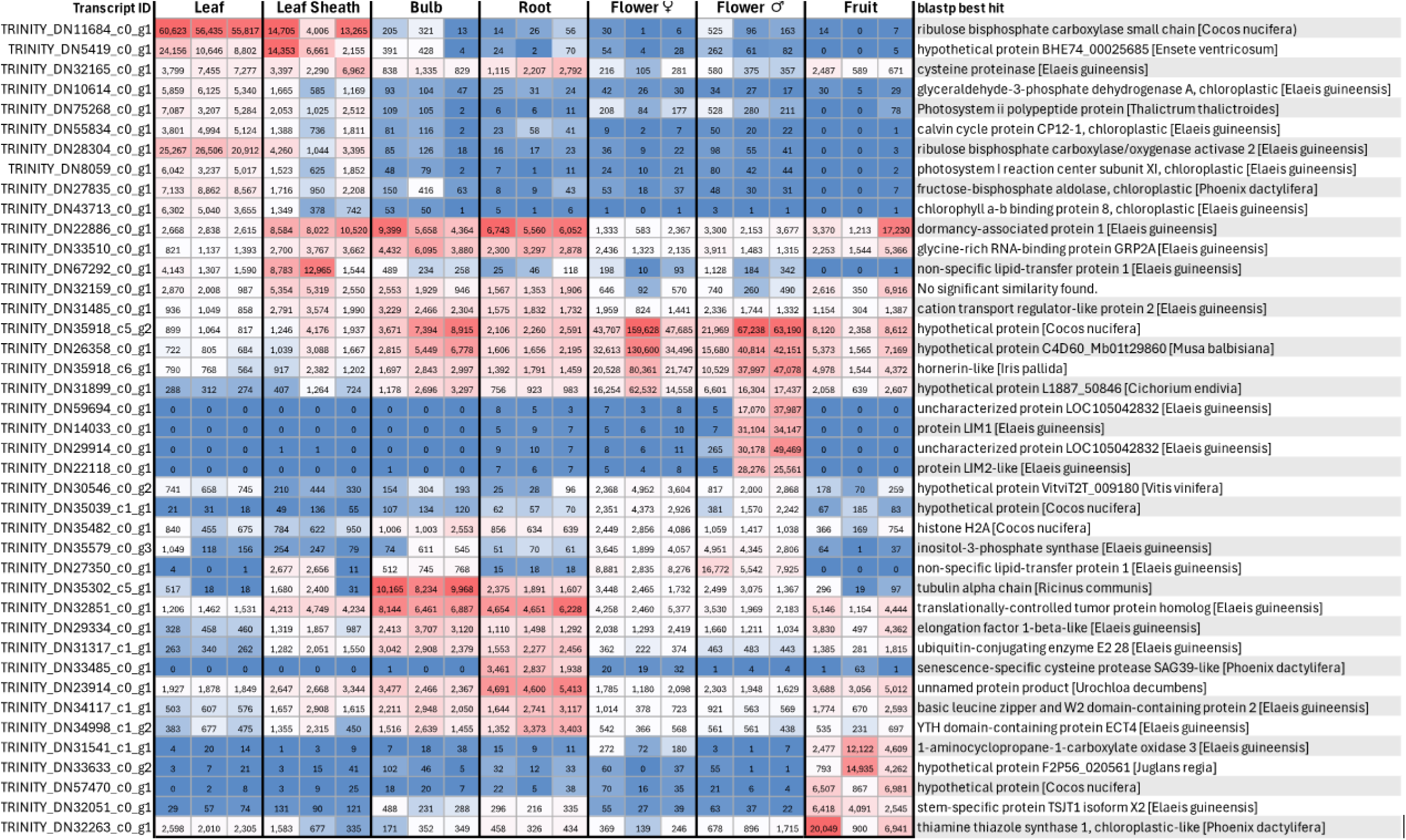
Heatmap of Transcripts-per-Million (TPM) values of the 10 transcripts with the highest median expression in each organ. The highest TPM values in each organ are presented in red and the lowest in blue.

In addition, a transcript annotated as translationally controlled tumor protein (TCTP) (TRINITY_DN32851_c0_g1) was found among the most highly expressed in the leaf sheaths, bulb, root, female flower, and fruit. TCTP is regulated at both translational and post-translational levels and is associated with various biochemical and cellular functions, including regulation of cell cycle progression, responses to abiotic and biotic stresses, plant growth, and signaling (Betsch et al., 2017; Wang et al., 2015). In fruits, one of the most highly expressed transcripts (TRINITY_DN31541_c1_g1), with a median TPM of 4,608.9, was identified as aminocyclopropanecarboxylate oxidase, an enzyme involved in ethylene production.

### 3.3 Organ-specific transcripts

A total of 306, 32, 41, 159, 158 and 916 transcripts were identified as organ-specific (SPM metric >= 0.95) in leaf, leaf sheaves, bulb, flowers, fruit and root, respectively (Supplementary File 2). Table 1 presents the top ten organ-specific transcripts, characterized by the highest tissue specificity (SPM – Specificity Metric Value) and the highest average gene expression levels (TPM – Transcripts Per Million) in each organ.

**Table 1.** The top ten organ-specific transcripts and the average gene expression (TPM – Transcripts Per Million) of each transcript. In the table: Transcript ID; Specificity metric value (SPM) for each transcript in each organ; Average of gene expression in TPM for each transcript; Organ specificity.

| Transcript ID | SPM | TPM | Organ-Specific |
| --- | --- | --- | --- |
| TRINITY_DN22255_c0_g1 | 0,9985 | 399,38 | Leaf Sheath |
| TRINITY_DN24567_c0_g1 | 0,9985 | 202,6433333 | Leaf Sheath |
| TRINITY_DN42235_c0_g1 | 0,9985 | 64,24666667 | Leaf Sheath |
| TRINITY_DN16207_c0_g1 | 0,9983 | 44,11 | Leaf Sheath |
| TRINITY_DN57848_c0_g1 | 0,9979 | 87,04333333 | Leaf Sheath |
| TRINITY_DN27363_c0_g1 | 0,997 | 280,2766667 | Leaf Sheath |
| TRINITY_DN42564_c0_g1 | 0,9936 | 8,206666667 | Leaf Sheath |
| TRINITY_DN25756_c0_g1 | 0,9902 | 90,29333333 | Leaf Sheath |
| TRINITY_DN27325_c0_g1 | 0,9892 | 49,93333333 | Leaf Sheath |
| TRINITY_DN29205_c0_g1 | 0,9885 | 208,4166667 | Leaf Sheath |
| TRINITY_DN32453_c0_g1 | 0,9979 | 197,7833333 | Leaf |
| TRINITY_DN18678_c0_g1 | 0,9978 | 87,52 | Leaf |
| TRINITY_DN25226_c0_g1 | 0,9976 | 64,39333333 | Leaf |
| TRINITY_DN42263_c0_g1 | 0,9967 | 351,66 | Leaf |
| TRINITY_DN12744_c0_g1 | 0,9965 | 334,86 | Leaf |
| TRINITY_DN6089_c0_g1 | 0,996 | 5501,643333 | Leaf |
| TRINITY_DN32895_c1_g1 | 0,9951 | 24,51333333 | Leaf |
| TRINITY_DN4338_c0_g1 | 0,9948 | 4045,5 | Leaf |
| TRINITY_DN29936_c0_g1 | 0,9945 | 817,5833333 | Leaf |
| TRINITY_DN81211_c0_g1 | 0,9942 | 183,6966667 | Leaf |
| TRINITY_DN64786_c0_g1 | 1 | 4843,65 | Fruit |
| TRINITY_DN57470_c0_g1 | 1 | 4784,81 | Fruit |
| TRINITY_DN25411_c0_g1 | 1 | 3745,27 | Fruit |
| TRINITY_DN30753_c1_g1 | 1 | 2472,01 | Fruit |
| TRINITY_DN83931_c0_g1 | 1 | 521,6366667 | Fruit |
| TRINITY_DN33633_c0_g2 | 0,9999 | 6663,42 | Fruit |
| TRINITY_DN68783_c0_g1 | 0,9999 | 111,7233333 | Fruit |
| TRINITY_DN16032_c0_g1 | 0,9993 | 395,3 | Fruit |
| TRINITY_DN31541_c1_g1 | 0,9992 | 6402,73 | Fruit |
| TRINITY_DN21620_c0_g1 | 0,9992 | 616,7766667 | Fruit |
| TRINITY_DN31555_c0_g1 | 0,997 | 177,1866667 | Bulb |
| TRINITY_DN17375_c0_g1 | 0,9965 | 177,9866667 | Bulb |
| TRINITY_DN35665_c0_g2 | 0,994 | 192,0733333 | Bulb |
| TRINITY_DN85352_c0_g1 | 0,9919 | 4,88 | Bulb |
| TRINITY_DN36994_c0_g1 | 0,9909 | 50,10333333 | Bulb |
| TRINITY_DN35321_c0_g1 | 0,9906 | 125,5233333 | Bulb |
| TRINITY_DN15982_c0_g1 | 0,9888 | 166,5666667 | Bulb |
| TRINITY_DN27700_c0_g1 | 0,9869 | 34,79 | Bulb |
| TRINITY_DN34950_c0_g1 | 0,9864 | 68,68 | Bulb |
| TRINITY_DN25927_c0_g2 | 0,9854 | 15,14333333 | Bulb |
| TRINITY_DN33485_c0_g1 | 1 | 2745,543333 | Root |
| TRINITY_DN35553_c0_g1 | 1 | 1359,476667 | Root |
| TRINITY_DN32252_c0_g1 | 1 | 759,4066667 | Root |
| TRINITY_DN32126_c0_g1 | 1 | 648,8333333 | Root |
| TRINITY_DN19440_c0_g1 | 1 | 512,7433333 | Root |
| TRINITY_DN31923_c0_g1 | 1 | 477,7 | Root |
| TRINITY_DN32518_c0_g2 | 1 | 450,0966667 | Root |
| TRINITY_DN29395_c0_g1 | 1 | 437,4666667 | Root |
| TRINITY_DN35639_c0_g1 | 1 | 434,2333333 | Root |
| TRINITY_DN20647_c0_g1 | 1 | 354,5833333 | Root |
| TRINITY_DN72992_c0_g1 | 0,9941 | 33,49666667 | F flower |
| TRINITY_DN82544_c0_g1 | 0,9926 | 97,26333333 | F flower |
| TRINITY_DN4600_c0_g1 | 0,9904 | 94,89 | F flower |
| TRINITY_DN51660_c0_g1 | 0,9899 | 26,39333333 | F flower |
| TRINITY_DN32222_c0_g1 | 0,985 | 2,793333333 | F flower |
| TRINITY_DN1988_c0_g1 | 0,9846 | 55,96333333 | F flower |
| TRINITY_DN33378_c0_g3 | 0,9843 | 20,01666667 | F flower |
| TRINITY_DN38420_c0_g1 | 0,9818 | 76,86333333 | F flower |
| TRINITY_DN28517_c0_g1 | 0,9809 | 216,98 | F flower |
| TRINITY_DN27824_c0_g1 | 0,9809 | 187,64 | F flower |
| TRINITY_DN29914_c0_g1 | 1 | 26637,36 | M flower |
| TRINITY_DN14033_c0_g1 | 1 | 21752,47333 | M flower |
| TRINITY_DN59694_c0_g1 | 1 | 18354,12667 | M flower |
| TRINITY_DN22118_c0_g1 | 1 | 17947,47667 | M flower |
| TRINITY_DN8432_c0_g1 | 1 | 1069,823333 | M flower |
| TRINITY_DN12894_c0_g1 | 0,9999 | 1045,283333 | M flower |
| TRINITY_DN28924_c0_g1 | 0,9995 | 154,58 | M flower |
| TRINITY_DN17272_c0_g1 | 0,9989 | 88,24 | M flower |
| TRINITY_DN65446_c0_g1 | 0,9986 | 50,32666667 | M flower |
| TRINITY_DN55061_c0_g1 | 0,9983 | 242,6966667 | M flower |

As expected, most leaf and leaf sheath-specific transcripts ere annotated as photosynthesis-related protein. In flowers, transcripts annotated as non-specific lipid-transfer protein (nsLTP), transcriptional regulator ICP4, histone proteins (mainly the H3 variant) and flavonoid reductase were identified. nsLTP is a well-known multigene family and its expression has been detected in the flowers of many plant species and may be associated with reproduction (Lee et al., 2009; Liu et al., 2015).

Interestingly, nine fruit-specific transcripts displayed higher TPM values compared to other organs and were annotated as enzymes involved in fatty acid synthesis and saturation, such as enoyl-[acyl-carrier-protein (ACP)] reductase – FabL (TRINITY_DN64786_c0_g1) and trans-2-enoyl-CoA reductase (TER), an enoyl-ACP reductase isoform – FabI (TRINITY_DN25411_c0_g1).

Root-specific transcripts included secretory peroxidases (18 transcripts), SGNH plant lipase (9 transcripts), phytocyanin (3 transcripts), Cytochrome P450 (35 transcripts), transcriptional regulator ICP4 (28 transcripts), and Serine/Threonine Kinases (29 transcripts (Supplementary File 2). The three most expressed root-specific transcripts were TRINITY_DN33485_c0_g1, TRINITY_DN25836_c0_g1 and TRINITY_DN35553_c0_g1, annotated as a cysteine peptidase (pfam00112), a phytocyanin (cd04216), and a peptidase S8 (cd04852), respectively. In addition, conserved domain associated with pathogen defense were identified in root-specific transcripts. These included crd07816, Bet_v1-like (TRINITY_DN32369_c0_g1), cd01960, non-specific lipid-transfer protein type 1 (nsLTP1) subfamily (TRINITY_DN47091_c0_g1), and cd02877, GH18_hevamine_XipI_class_III (TRINITY_DN34687_c0_g1). These protein families play a central role in plant defense against a broad spectrum of biotic and abiotic stresses, particularly in those caused by fungal and bacterial defense (Sudisha et al., 2012; Van Loon and Van Strien, 1999).

### 3.4 Expanded gene families in the macaúba palm

Gene family analysis was performed using proteome data of *Elaeis guineensis* (African oil palm)*, Musa acuminata* (banana)*, Phoenix dactylifera* (date palm) and *Oryza sativa* (rice) genomes, and the transcriptome of macaúba palm. A total of 15,283 orthogroups (at least one species representing the group) were identified and 12,664 family groups have one representant of macaúba palm. Phylogenetic analysis was undertaken using only single copy gene families (Supplementary figure 5). The maximum likelihood bootstrap values are shown as percentage.

Macaúba palm and African oil palm were first clustered into the same clade, with date palm as a sister clade, establishing a palm tree cluster. That data is consistent since African oil palm and macaúba palm are from the same Tribe (Tribe Cocoseae; Family Arecoideae) and Date palm belongs to another subfamily (Coryphoideae). All the palm trees (Arecaceae) belong to a monophyletic group including five subfamilies, 28 tribes and 27 subtribes (Baker and Dransfield, 2016). Despite a reduced number of palm tree samples in our data, the monophyly of the Arecaceae family is supported by the 100% of bootstraps.

Gene family expansion analysis revealed 20 expanded gene families in macaúba palm specie compared with African oil palm (Table 2). Among these, families were associated with stress responses and developmental regulation, including the Pentatricopeptide repeat-containing protein family (PPR) (OG0000057, OG0000748, OG0002240, OG0002627), leucine-rich repeat receptor-like protein kinase (LRR-RLK) (family OG0000697), serine/threonine-protein kinase (OG0001613 and OG0002773), and putative disease resistance protein RGA (OG0003836) from the same group of PPR and RLK proteins. These protein families are associated with the main function in pathogen recognition and activation of defense pathways.

**Table 2.**
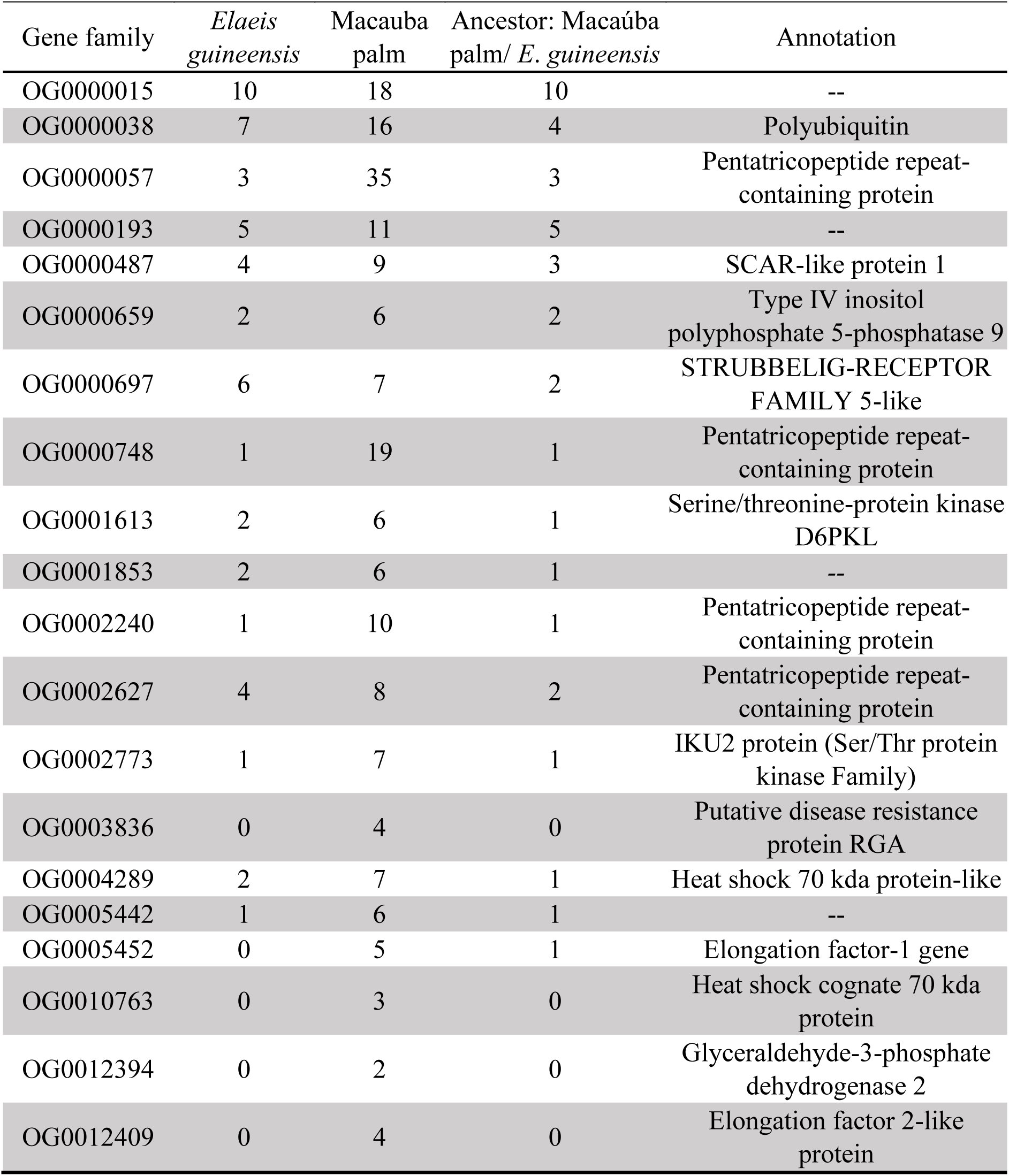
Gene families expanded in macaúba palm compared to African Oil Palm (*Elaeis guineensis*) and a common ancestor. In the table: Gene family name; the number of African Oil Palm genes; number of macaúba palm transcripts; number of common ancestor genes; and functional annotation; The false discovery rate (FDR) for all families were FDR < 0.05.

In family OG0001613, four transcripts with high TPM values (TRINITY_DN11200_c0_g1, TRINITY_DN25915_c0_g1, TRINITY_DN34203_c0_g1, TRINITY_DN34203_c1_g3) were like D6PKL (D6 PROTEIN KINASE-LIKE). D6PKL is a kinase that regulates the auxin transport activity of PIN auxin efflux facilitators by direct phosphorylation. D6PK-mediated PIN phosphorylation promotes auxin transport in the hypocotyl, which is a prerequisite for PHOT1-dependent hypocotyl bending. PHOT1 (phototropin 1) regulates light responses, including stomatal opening, thereby controlling CO₂ uptake for photosynthesis and water loss through transpiration (Kinoshita et al., 2001; Weller et al., 2017; Willige et al., 2013). Additionally, 55 gene families were exclusively found in macaúba palm but were absent from the other palm species analyzed (*Elaeis guineensis* and *Phoenix dactylifera*). Of these, half of them (28) are hypothetical proteins (Supplementary File 3), including two transcripts (OG0014852: TRINITY_DN26358_c0_g1 and OG0015050: TRINITY_DN35918_c6_g1) encoding for hypothetical proteins that were highly expressed in both male and female flowers (Figure 3). Furthermore, a histone H4 gene (TRINITY_DN34622_c0_g1), closely related to fern species, was identified as highly expressed in female flowers. Another gene, encoding a ubiquitin protein (TRINITY_DN33606_c3_g2), exhibited phylogenetic similarity to eudicot species and was expressed across all organs, with particularly high expression in leaf, bulb, and root.

A total of 48 gene families were shared between macaúba and date palm, whereas 85 were shared between macaúba and African oil palm (Supplementary File 3). Table 3 presents the median TPM values of the shared families that were annotated with KEGG Orthology (KO). The gene family OG0014105, shared with date palm, exhibited the highest TPM values and was annotated as a protein phosphatase 2C (PP2C), with its highest expression observed in the root and leaf sheath. Meanwhile, gene family OG0006978, shared with African oil palm, had the highest TPM values and was annotated as EMC8_9, an ER membrane protein complex subunit 8/9, showing peak expression in the bulb, root, and fruit.

**Table 3.** Overview of gene families shared between macaúba palm, date palm, and African oil palm. This table presents general information on the gene families common to macaúba palm and date palm, as well as those shared between macaúba palm and African oil palm. The median Transcripts-per-Million (TPM) values for each gene family in different organs are displayed. Cell highlighting intensity corresponds to the expression levels of each transcript, with darker shades indicating higher expression of each transcript. The KEGG orthology functions and pathways associated with each gene family are also provided.

| Gene family | Shared with | Median TPM |  |  |  |  |  |  | Function | Pathway |
| --- | --- | --- | --- | --- | --- | --- | --- | --- | --- | --- |
|  |  | Leaf | Leaf Sheath | Bulb | Root | F flower | M flower | Fruit |  |  |
| OG0013975 | Date palm | 0 | 0 | 0 | 2.87 | 0.05 | 0.18 | 0 | E1.11.1.7; peroxidase | Phenylpropanoid biosynthesis [PATH:ko00940] |
| OG0014810 | Date palm | 1.77 | 1.57 | 4.18 | 5.38 | 4.38 | 3.48 | 0.65 | MTERF; transcription termination factor, mitochondrial | Translation factors [BR:ko03012] |
| OG0014851 | Date palm | 0.29 | 0.16 | 0.43 | 6.03 | 0.77 | 0.4 | 1.4 | SAUR; SAUR family protein | Plant hormone signal transduction [PATH:ko04075] |
| OG0014918 | Date palm | 2.45 | 1.92 | 3.7 | 3.85 | 1.92 | 3.18 | 2.67 | DGAT1; diacylglycerol O-acyltransferase 1 | Glycerolipid metabolism [PATH:ko00561] |
| OG0006474 | Date palm | 33.65 | 30.79 | 26.09 | 34.49 | 25.24 | 26.75 | 15.85 | alpha- and gamma-adaptin-binding protein p34 | Membrane trafficking [BR:ko04131] |
| OG0014105 | Date palm | 16.26 | 41.11 | 17.92 | 47.28 | 6.05 | 7.11 | 8.32 | PP2C; protein phosphatase 2C | MAPK signaling pathway - plant [PATH:ko04016] |
| OG0014073 | Date palm | 0.04 | 0 | 0 | 7.13 | 0.02 | 22.89 | 0.37 | E3.2.1.4; endoglucanase | Starch and sucrose metabolism [PATH:ko00500] |
| OG0010775 | Date palm | 9.37 | 8.66 | 11.86 | 12.91 | 10.31 | 17.11 | 17.49 | SDHAF1; succinate dehydrogenase assembly factor 1 | Mitochondrial biogenesis [BR:ko03029] |
| OG0014061 | Date palm | 0.14 | 4.66 | 3.48 | 2.74 | 19.91 | 22.8 | 3.89 | COX2; cytochrome c oxidase subunit 2 | Oxidative phosphorylation [PATH:ko00190] |
| OG0014104 | Date palm | 0.59 | 11.17 | 8.02 | 6.28 | 23.55 | 22.71 | 0.94 | ND4; NADH-ubiquinone oxidoreductase chain 4 | Oxidative phosphorylation [PATH:ko00190] |
| OG0014891 | Date palm | 1.75 | 2.26 | 3.87 | 3.46 | 22.28 | 21.57 | 0.77 | RP-S13, rpsM; small subunit ribosomal protein S13 | Ribosome [PATH:ko03010] |
| OG0008562 | Oil palm | 13.66 | 17.2 | 0.78 | 11.39 | 0.39 | 0.46 | 0.03 | MFS transporter, POT/PTR family, nitrate transporter | Transporters [BR:ko02000] |
| OG0009269 | Oil palm | 7.71 | 3.19 | 4.24 | 7.62 | 0.02 | 0 | 5.74 | AOC3, AOC2, tynA; primary-amine oxidase | Glycine, serine and threonine metabolism [PATH:ko00260] |
| OG0008552 | Oil palm | 7.12 | 13.31 | 13.71 | 20.38 | 15.12 | 14.68 | 13.03 | ATG16L1; autophagy-related protein 16-1 | Autophagy - animal [PATH:ko04140] |
| OG0013626 | Oil palm | 0 | 0.02 | 0.13 | 43.85 | 0.33 | 0.29 | 0.02 | PRUNE, PPX1; exopolyphosphatase | Purine metabolism [PATH:ko00230] |
| OG0014241 | Oil palm | 0.72 | 0.87 | 9.05 | 15.45 | 0.23 | 0.59 | 3.21 | GST, gst; glutathione S-transferase | Glutathione metabolism<br>[PATH:ko00480] |
| OG0006978 | Oil palm | 44.2 | 103.11 | 150.59 | 148.52 | 92.57 | 112.13 | 128.98 | EMC8_9; ER membrane protein complex subunit 8/9 | Membrane trafficking<br>[BR:ko04131] |
| OG0012991 | Oil palm | 12.12 | 16.43 | 25.48 | 30.19 | 19.68 | 21.17 | 23.21 | EIF2B5; translation initiation factor eIF-2B subunit epsilon | RNA transport<br>[PATH:ko03013] |
| OG0011015 | Oil palm | 28.26 | 31.45 | 30.54 | 35.68 | 33.42 | 23.55 | 32.58 | UTP7, WDR46; U3 small nucleolar RNA-associated protein 7 | Ribosome biogenesis<br>[BR:ko03009] |
| OG0012902 | Oil palm | 12.81 | 23.86 | 27.31 | 28.66 | 19.58 | 11.97 | 13.68 | GEMIN2, SIP1; gem associated protein 2 | RNA transport<br>[PATH:ko03013] |
| OG0013101 | Oil palm | 11.83 | 17.5 | 22.5 | 22.67 | 16.16 | 14.03 | 8.14 | VPS11, PEP5; vacuolar protein sorting-associated protein 11 | Autophagy - yeast<br>[PATH:ko04138] |
| OG0009566 | Oil palm | 57.44 | 33.76 | 34.91 | 40.66 | 37.43 | 40.84 | 23.36 | MTR3, EXOSC6; exosome complex component MTR3 | RNA degradation<br>[PATH:ko03018] |
| OG0013182 | Oil palm | 151.96 | 69.86 | 36.2 | 21.91 | 27.63 | 24.73 | 36.3 | RP-L4, MRPL4, rplD; large subunit ribosomal protein L4 | Ribosome<br>[PATH:ko03010] |
| OG0006579 | Oil palm | 83.35 | 28.06 | 15.6 | 10.41 | 19.62 | 31.31 | 22.5 | SRX1; sulfiredoxin | Enzymes with EC numbers |
| OG0014651 | Oil palm | 10.05 | 13.1 | 3.9 | 4.78 | 2.5 | 0.83 | 0.14 | TOGT1; scopoletin glucosyltransferase | Phenylpropanoid biosynthesis<br>[PATH:ko00940] |
| OG0011540 | Oil palm | 104.35 | 73.57 | 45.49 | 43.38 | 44.74 | 42.91 | 78.23 | ENY2, DC6, SUS1; enhancer of yellow 2 transcription factor | Transcription machinery<br>[BR:ko03021] |
| OG0014242 | Oil palm | 9.03 | 13.52 | 17.07 | 18.42 | 10.09 | 9.58 | 21.97 | FBXO7; F-box protein 7 | Ubiquitin system<br>[BR:ko04121] |
| OG0010940 | Oil palm | 11.33 | 20.11 | 14.98 | 17.16 | 5.39 | 13.8 | 38.09 | RHBDD1; rhomboid domain-containing protein 1 | Peptidases<br>[BR:ko01002] |
| OG0011418 | Oil palm | 42.23 | 66.9 | 79.88 | 63.49 | 79.89 | 33.15 | 48.91 | C1QBP; complement component 1 Q subcomponent-binding protein, mitochondrial | Glycosaminoglycan binding proteins<br>[BR:ko00536] |
| OG0004673 | Oil palm | 21.37 | 64.76 | 59.14 | 46.11 | 88.32 | 56.29 | 13.24 | SDHC, SDH3; succinate dehydrogenase (ubiquinone) cytochrome b560 subunit | Citrate cycle (TCA cycle)<br>[PATH:ko00020] |
| OG0012970 | Oil palm | 1.85 | 1.8 | 4.72 | 3.42 | 12.97 | 7.73 | 1.84 | MCM10; minichromosome maintenance protein 10 | DNA replication proteins<br>[BR:ko03032] |
| OG0013109 | Oil palm | 30.84 | 55.88 | 111.94 | 83.2 | 152.36 | 70.31 | 63.93 | RPB3, POLR2C; DNA-directed RNA polymerase II subunit RPB3 | RNA polymerase<br>[PATH:ko03020] |

### 3.5 Transcripts related to hydric stress

We analyzed gene expression related to drought stress processes by mapping macaúba palm transcripts to orthologs from *Arabidopsis thaliana, Zea mays*, *Oryza sativa*, and *Elaeis guineensis* using the Drought DB – Drought Stress Gene Database (Alter et al., 2015). The putative orthologs were grouped into gene families. A total of 221 macaúba palm transcripts were identified across 112 families, each containing at least one transcript (Supplementary File 4).

Transcription factors (TFs) are well-known, playing a key role in regulating gene expression and are relatively conserved across bacteria, humans, and plants. Among several families identified, transcription factors associated with drought stress response exhibited high expression levels across all organs analyzed. These included transcription factors MYC (family OG0000052), bZIP (families OG0000057 and OG0000069), MYB (OG0000315 and OG0000932), NF-Y (OG0000005 and OG0000012), ERF-ethylene-responsive transcription factor (OG0000796, OG0000112, OG0000209), AP2 (OG0001372), WRKY (OG0002791), and HSFA1 (family OG0000123).

The MYC, bZIP, and MYB transcription factors are known to be regulated by ABA (abscisic acid), a key hormone involved in plant stresses response activating signaling cascades triggered by dehydration signal. In our dataset, transcripts TRINITY_DN2149_c0_g1, TRINITY_DN27638_c0_g2, and TRINITY_DN31421_c1_g1 from MYC family (OG0000052) exhibited high expression across all tissues. For the MYB families (OG0000932 and OG0000315) only transcript TRINITY_DN33427_c1_g2 displayed high expression, (TPM ≥ 20), particularly in the vegetative tissues and fruit.

The MYC transcription factor has previously been reported to confer cold tolerance when a maize MYC-type ICE-like transcription factor is ectopically expressed in *Arabidopsis thaliana*. Both MYC and MYB act as transcriptional activators of ABA-inducible gene expression under drought stress in *Arabidopsis thaliana* (Abe et al., 2003; Lu et al., 2017). Moreover, bZIP families OG0000057 and OG0000069 also showed high expression in all analyzed organs. In OG0000057, TRINITY_DN28492_c0_g1 and TRINITY_DN63679_c0_g1 were highly expressed in all organs, while TRINITY_DN21593_c0_g1 was not expressed in leaf tissue. In OG0000069 family, TRINITY_DN22238_c0_g1 and TRINITY_DN28034_c0_g1 were expressed in leaf, leaf sheave, bulb, and root organs.

The AP2/ERF (APETALA2/ethylene responsive factor) superfamily, one of the largest groups of transcription factors in plants, was also well represented. ERF (ethylene-responsive factor) family transcripts were distributed into the families OG0000796, OG0000112, OG0000209. Among these, only OG0000112 displayed high expression, with transcript TRINITY_DN33317_c0_g1 expressed specifically in the fruit. Additionally, TRINITY_DN30070_c0_g1, from AP2/ERF family OG0001372, was highly expressed in root, as found in sesame (Dossa et al., 2016), and male flower.

Of the macaúba palm transcripts with DroughtDB orthologs, 20 transcripts (in 13 families) showed organ-specific expression (Table 4). Notably, two transcripts (TRINITY_DN26933_c0_g1 and TRINITY_DN35293_c0_g1) from transcription repressor MYB5-like family (OG0000065) were specific to the root, while TRINITY_DN22101_c0_g1 showed specificity to the leaf sheath. Similarly, one transcript from OG0000034 (respiratory burst oxidase homolog protein E isoform X2) was specific to the root (TRINITY_DN24271_c0_g1) and another to the fruit (TRINITY_DN35726_c1_g1).

**Table 4.** Overview of organ-specific transcripts with orthologous of drought tolerance. Transcripts with orthologous genes found in the DroughtDB and SPM (specificity measure) >=0.95.

| Transcript ID | Gene family | Median TPM |  |  |  |  |  |  | Specific | Function |
| --- | --- | --- | --- | --- | --- | --- | --- | --- | --- | --- |
|  |  | Leaf | Leaf Sheath | Bulb | Root | F flower | M flower | Fruit |  |  |
| TRINITY_DN82979_c0_g1 | OG0000003 | 0.46 | 0.9 | 2.53 | 4.4 | 2.26 | 18.95 | 1.67 | Flower M | calcium-dependent protein kinase 26 |
| TRINITY_DN9444_c0_g1 | OG0000008 | 2.17 | 6.02 | 1.38 | 34.71 | 0.47 | 1.89 | 0.05 | Root | beta-glucosidase 12-like |
| TRINITY_DN22700_c0_g1 | OG0000018 | 0.26 | 1.63 | 5.27 | 19.02 | 0.08 | 0.26 | 0.05 | Root | ABC transporter G family member 45-like isoform X1 |
| TRINITY_DN31780_c2_g1 | OG0000018 | 1.86 | 3.18 | 7.5 | 42.69 | 1.18 | 1.62 | 0.91 | Root | ABC transporter G family member 45-like isoform X1 |
| TRINITY_DN31836_c2_g1 | OG0000018 | 0.29 | 0.46 | 0.46 | 5.42 | 0.36 | 0.27 | 0.08 | Root | ABC transporter G family member 45-like isoform X1 |
| TRINITY_DN18901_c0_g1 | OG0000027 | 0.2 | 0.36 | 0.2 | 4.69 | 0.2 | 0.47 | 0.37 | Root | plasma membrane ATPase-like |
| TRINITY_DN34220_c0_g1 | OG0000027 | 8.05 | 10.23 | 7.94 | 90.01 | 8.31 | 13.25 | 12.67 | Root | plasma membrane ATPase-like |
| TRINITY_DN24271_c0_g1 | OG0000034 | 0 | 0.18 | 1.63 | 9.34 | 0.04 | 0 | 0.21 | Root | respiratory burst oxidase homolog protein E isoform X2 |
| TRINITY_DN35726_c1_g1 | OG0000034 | 3.08 | 8.77 | 21.06 | 22.17 | 6.9 | 14.04 | 174.54 | Fruit | respiratory burst oxidase homolog protein E isoform X2 |
| TRINITY_DN22101_c0_g1 | OG0000065 | 0.1 | 12.89 | 1.71 | 0.91 | 1.39 | 1.98 | 0.22 | Leaf Sheath | transcription repressor MYB5-like |
| TRINITY_DN26933_c0_g1 | OG0000065 | 0 | 0.11 | 1.57 | 7.19 | 0 | 0 | 0 | Root | transcription repressor MYB5-like |
| TRINITY_DN35293_c0_g1 | OG0000065 | 0 | 0.52 | 4.88 | 16.7 | 0 | 0 | 0 | Root | transcription repressor MYB5-like |
| TRINITY_DN33317_c0_g1 | OG0000112 | 1.33 | 13.3 | 22.82 | 9.82 | 12.77 | 10.62 | 309.73 | Fruit | ethylene-responsive transcription factor ERF003-like |
| TRINITY_DN80402_c0_g1 | OG0000209 | 2.84 | 24.6 | 2.01 | 3.64 | 0.42 | 0 | 1.08 | Leaf Sheath | ethylene-responsive transcription factor ERF027-like |
| TRINITY_DN19269_c1_g1 | OG0000213 | 0.02 | 0 | 0.04 | 18.22 | 0 | 0 | 0 | Root | G-type lectin S-receptor-like serine/threonine-protein kinaseAT5g24080 |
| TRINITY_DN32893_c0_g2 | OG0000213 | 0.1 | 0.52 | 1.22 | 11.25 | 0.04 | 0.05 | 0.05 | Root | G-type lectin S-receptor-like serine/threonine-protein kinaseAT5g24080 |
| TRINITY_DN62888_c0_g1 | OG0000226 | 4.06 | 10.46 | 10.92 | 27.06 | 1.46 | 0.95 | 170.29 | Fruit | glutathione S-transferase U17-like |
| TRINITY_DN28160_c0_g1 | OG0000315 | 0.43 | 2.21 | 1.19 | 12.97 | 0.05 | 0 | 2.66 | Root | transcription factor MYB108-like |
| TRINITY_DN27408_c1_g1 | OG0001311 | 31.62 | 73.13 | 80.09 | 326.7 | 42.02 | 33.38 | 1735.5 | Fruit | abscisic stress-ripening protein 2-like |
| TRINITY_DN27479_c0_g1 | OG0002008 | 10 | 7.84 | 2.08 | 100.85 | 1.21 | 2.62 | 0.12 | Root | glu S.griseus protease inhibitor-like |

### 3.6 Fungi transcripts in the root

Surprisingly, we identified 3.9% of the root reads and a total of 7.091 assembled transcripts assigned as fungi (Figure 4). Most of these genes (86.6%) were assigned to arbuscular mycorrhizal fungi (AMF) *Rhizophagus* spp. (Glomerales), indicating endophytic symbiotic relationship between macaúba palm roots and AMF. Among these fungal transcripts, TRINITY_DN30513_c0_g1 encodes a 301 aa secreted protein (Sec/SPI, probability = 0.9991) of unknown function showed the highest-level expression (Supplementary File 5). The second most expressed fungal transcript TRINITY_DN33652_c0_g1 encodes for a 420 aa fatty-acid desaturase that showed 92% of identity to the OLE1-like protein of *Rhizophagus irregularis* (PKY45226.1).

**Figure 4.**
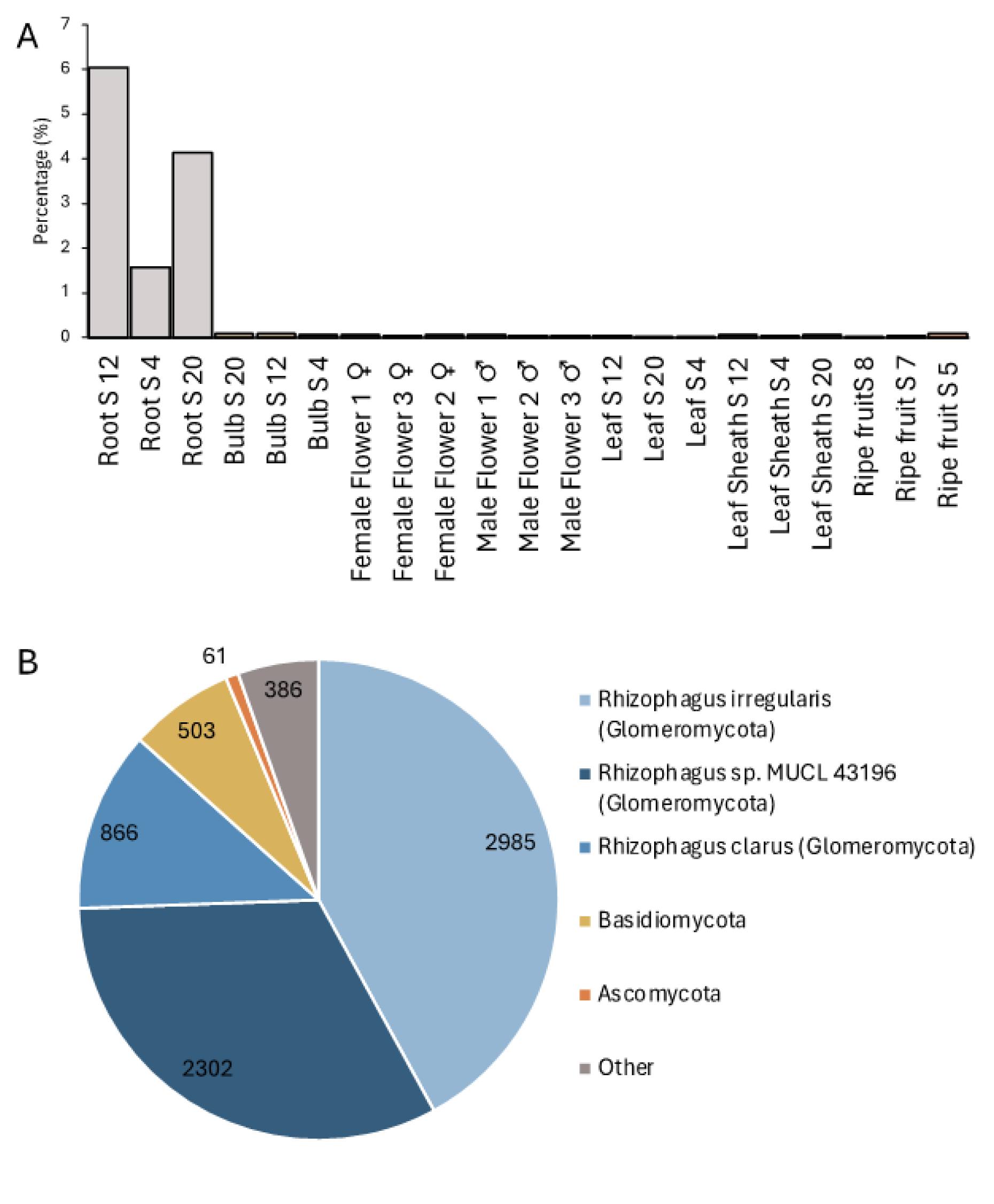
A. Percentage of reads identified as fungi in each sample. B. Taxonomic classification of the fungal transcripts.

## 4. Discussion

Morphological and structural adaptations are associated with biochemical and molecular changes that allow the species survival in several environments. In the generated data, first comprehensive transcriptomic atlas provides a valuable genomic resource to further research on evolutionary biology, genotype–phenotype associations, and molecular mechanistic studies. An extensive transcriptome characterization of multiple organs, and the annotation of transcripts allowed us to identify various predicted genes involved in key biochemical processes for both abiotic and biotic stress responses.

Organ-specific gene expression patterns underscore the specialized biological processes occurring in each organ (Birnbaum et al., 2003). For instance, the distinct upregulation of genes involved to photosynthesis in leaves and leaf sheaths, as well as the upregulation of nitrogen metabolism genes in roots, emphasizes the critical role of these organs in energy production and nutrient uptake, respectively. The presence of transcripts encoding proteins of unknown functions in reproductive organs suggests the existence of uncharacterized pathways important for the development of floral organs. The high expression of fatty acid synthases in fruit pulp supports a metabolic focus on oil accumulation, a trait of significant interest for biofuel production (Wong et al., 2017). Meanwhile, the enrichment of defense-related genes in roots, including those associated with pathogen defense, emphasizes the critical role of this organ in maintaining plant health under biotic stress (Van Loon and Van Strien, 1999).

The expansion of gene families, such as Pentatricopeptide repeat-containing proteins (PPR) and leucine-rich repeat receptor-like protein kinases (LRR-RLK), may contribute to the resilience of macaúba to diverse environmental conditions (Liu et al., 2017). Notably, the identification of 55 gene families unique to macaúba palm suggests that this species contains unique genetic elements that could be exploited for crop improvement.

The drought stress-related transcript analysis provides a deeper understanding of how macaúba palm may cope with hydric stress. The high expression of transcription factors AP2/ERF, MYC, bZIP, and MYB across various organs, along with their established roles in ABA-mediated stress responses, suggest that these factors play a central role in the drought tolerance (Jin et al. 2018). The organ-specific expression of certain drought-related genes further demonstrate organ-specific adaptation strategies, particularly in roots, which are essential for water uptake and retention (Gupta et al. 2020).

To our knowledge this is the first study to report an association between the AMF genus *Rhizophagus* spp. and macaúba palm. Although symbiotic associations with AMF have been reported for economically important palm species, including oil palm, date palm, and coconut (da Silva Maia et al., 2021; El Hilali et al., 2022; Lara-Pérez et al., 2020; Gagou et al. 2023), the association with macaúba palm remains poorly understood. The identification of a significant number of AMF transcripts in roots supports the existence of symbiotic relationship, through which improved nutrient uptake and increased tolerance to abiotic stresses.

*Rhizophagus irregularis*, a well-known AMF, has been shown to confer plant tolerance to abiotic stress conditions like drought and salinity by improving water uptake, nutrient acquisition, and activating antioxidant defense systems (Zhang et al, 2024; Liu et al, 2023; Alam et al., 2023). Of AMF-related transcripts, one of the highest expressed identified in our study, OLE1-like protein, is involved in the synthesis of palmitvaccenic acid and is conserved among AMF (order Glomerales) but not present in nonsymbiotic Mucoromycota. Palmitvaccenic acid (C16:1^Δ11cis^), a fatty acid synthesized by *Rhizophagus irregularis* and other AMF, is considered a lipid biomarker for AMF. Importantly, palmitvaccenic acid has been implicated in the establishment of AMF-plant symbiosis, as it can induce hyphal branching and spore formation in this species (Brands et al., 2020; Kameoka and Gutjahr, 2022).

The macaúba palm (*Acrocomia aculeata*) exhibit morphological and phenotypical adaptations that enable its survival across a wide range of edaphoclimatic conditions, mainly in semi-arid regions. According to Osório et al. 2013, species with drought tolerance mechanisms often present high phenotypic plasticity, a characteristic already described for the macaúba palm (Colombo et al., 2018; Pires et al., 2013; Marques et al., 2017). Among its phenotypic adaptations, the presence of leaflets covered with cuticle, deposits of epicuticular wax, abaxial stomata distribution and the protection of stipe by petioles bases are traits that contribute to its resilience under drought, high temperatures and fire (Vianna et al., 2017, Colombo et al., 2018, Bicalho et al., 2016).

The genome and transcriptome data provide invaluable insights about plasticity and rusticity of macaúba palm, as well as the differences compared to African oil palm. These findings are relevant considering the potential for oil production, and its wide range of applications and products. Macaúba is also considered a sustainable alternative in the context of energy transition, particularly for production of renewable diesel and SAF (sustainable aviation fuel). The first comprehensive overview of the transcriptomic landscape of macaúba palm highlight the molecular basis of organ-specific functions, stress responses, and symbiotic interactions. Although the data represent a single time point - limiting conclusions about physiology process, and metabolism of macaúba palm - it represents offers significant resource for a range of applications, including breeding, and functional genomics for improving oil yield and stress tolerance in macaúba palm.

## Supporting information

Macauba palm transcripts sequences.

SPM values per tissue, indicative of organ-specificity.

Gene family comparison in macauba palm, African oil palm and date palm.

Gene families containing at least one macauba palm transcript related to drought stress.

TPM values of transcripts assigned as fungi.

## 5. Acknowledgements

We would like to express our sincere gratitude to the Center for Computational Engineering and Sciences (CCES), Acelen Renováveis—through its Agribusiness Division under the framework of a scientific cooperation agreement with the University of Campinas (UNICAMP)—and Agronomic Institute of São Paulo, for their financial support and access to high-performance computing resources, which were essential to the success of this research project. This partnership enabled the development of the present study on gene expression in Acrocomia aculeata and its relevance to sustainable bioenergy applications.

