## Supplementary figures. for "A Transcriptomic Atlas of Macaúba Palm Reveals Organ-Specific Gene Expression and Stress-Related Pathways"

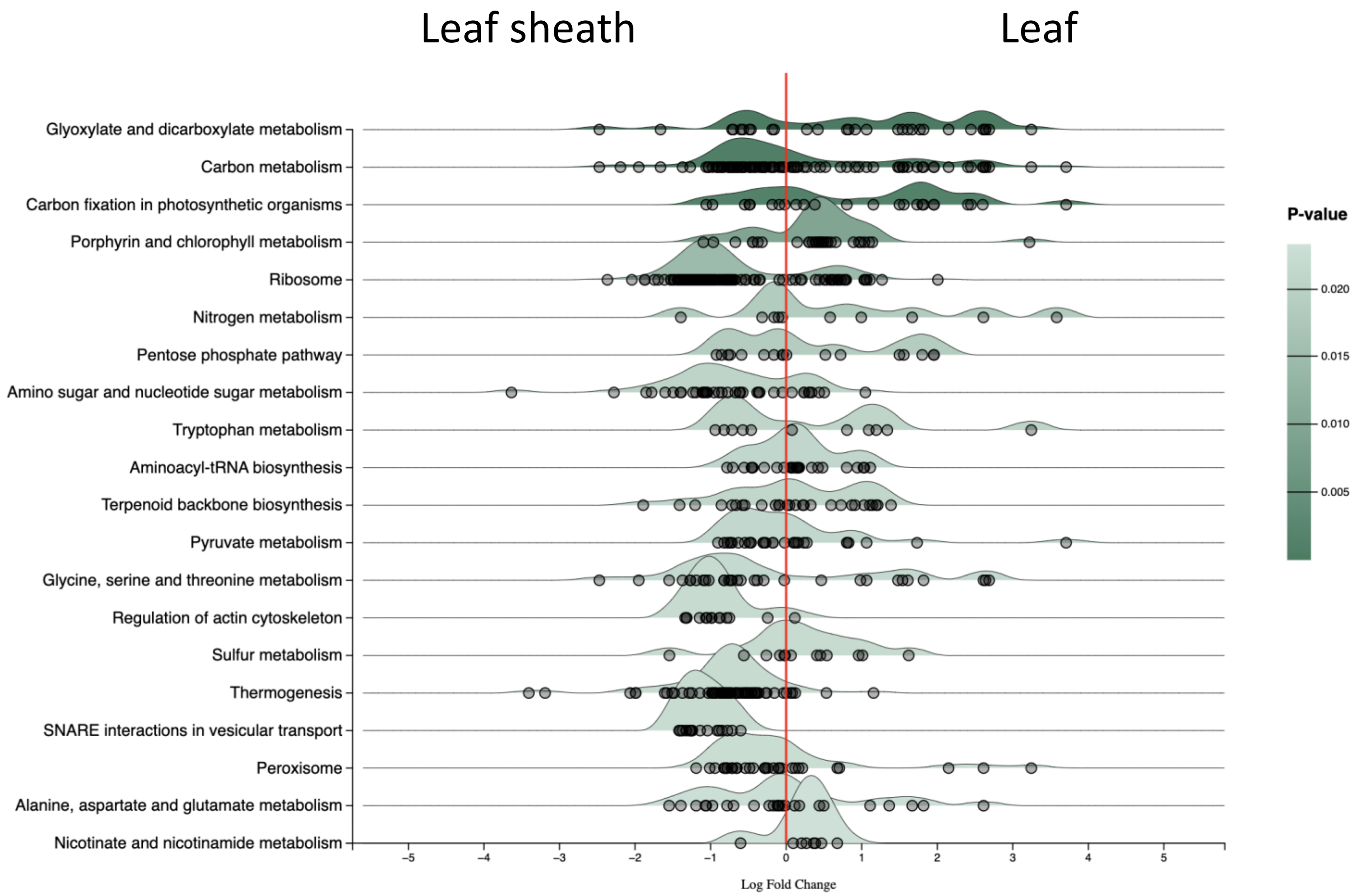


**Supplementary Figure 1.** Ridge plot of KEGG pathways defined by the Over-representation analysis of the transcriptome of leaf sheath and leaf. The circles represent genes in each pathways that differentially expressed between the organs. Pathways are shown in decreasing order of log2 fold change.

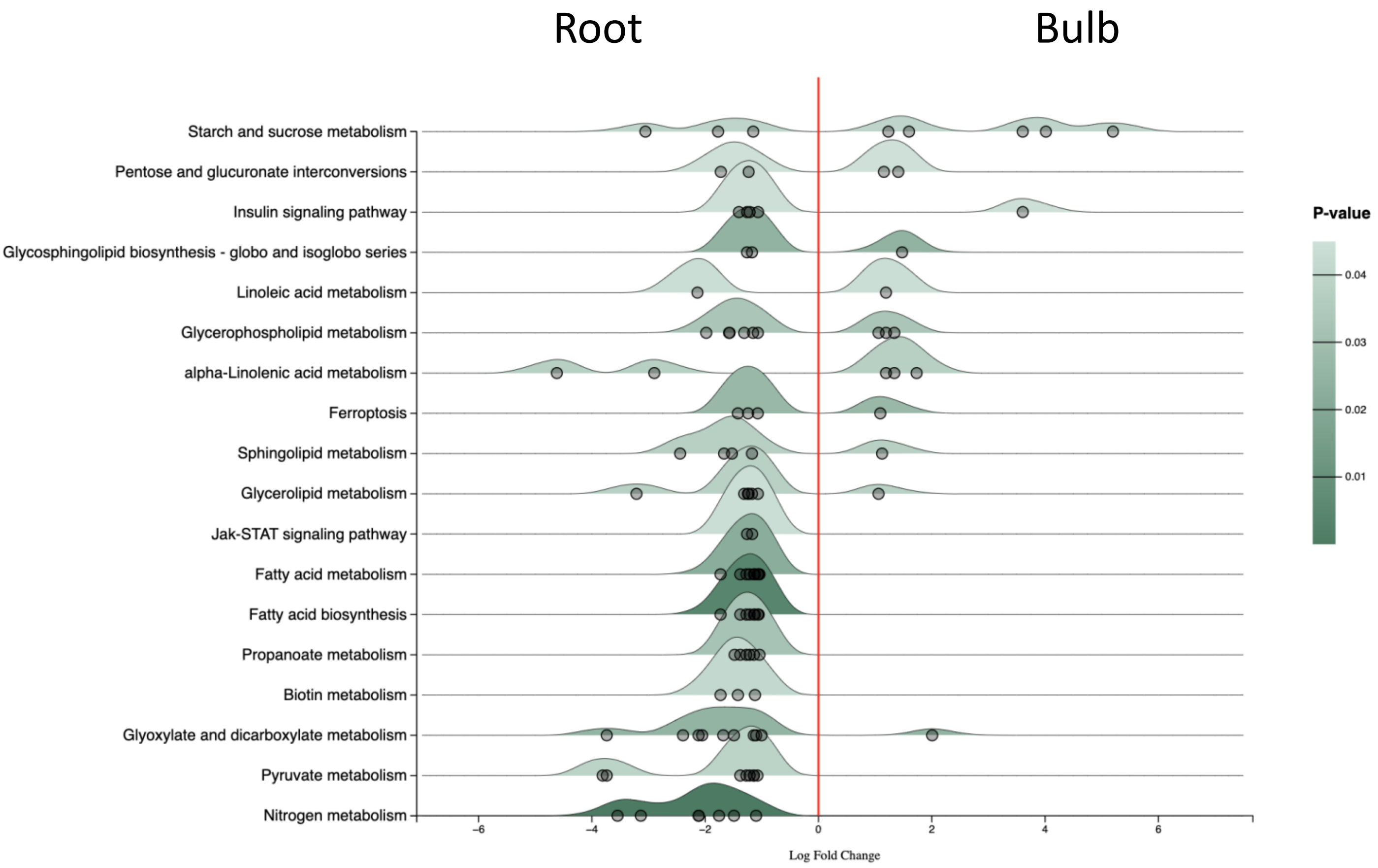


**Supplementary Figure 2.** Ridge plot of KEGG pathways defined by the Over-representation analysis of the transcriptome of root and bulb. The circles represent genes in each pathways that differentially expressed between the organs. Pathways are shown in decreasing order of log2 fold change.

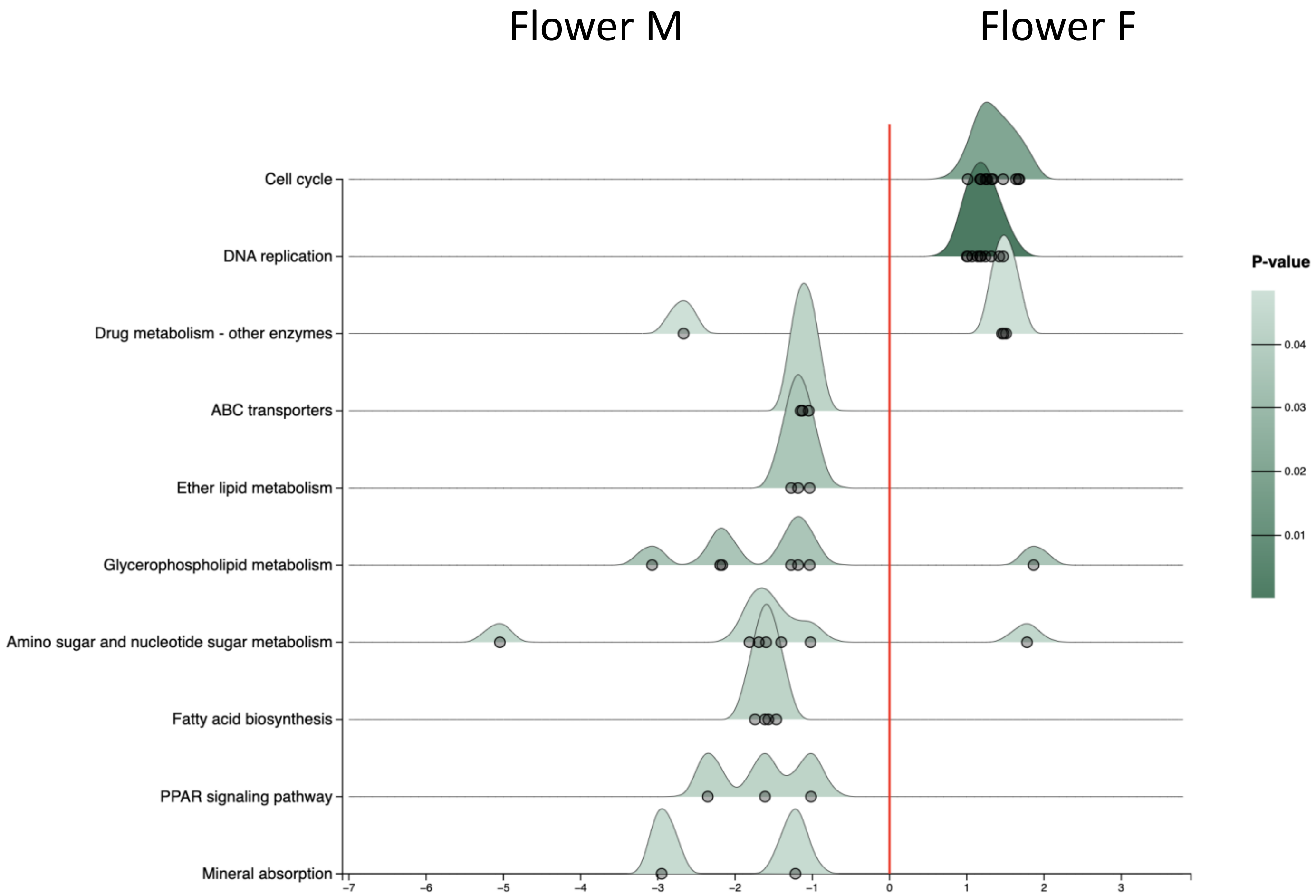


**Supplementary Figure 3.** Ridge plot of KEGG pathways defined by the Over-representation analysis of the transcriptome of male flower and female flower. The circles represent genes in each pathways that differentially expressed between the organs. Pathways are shown in decreasing order of log2 fold change.

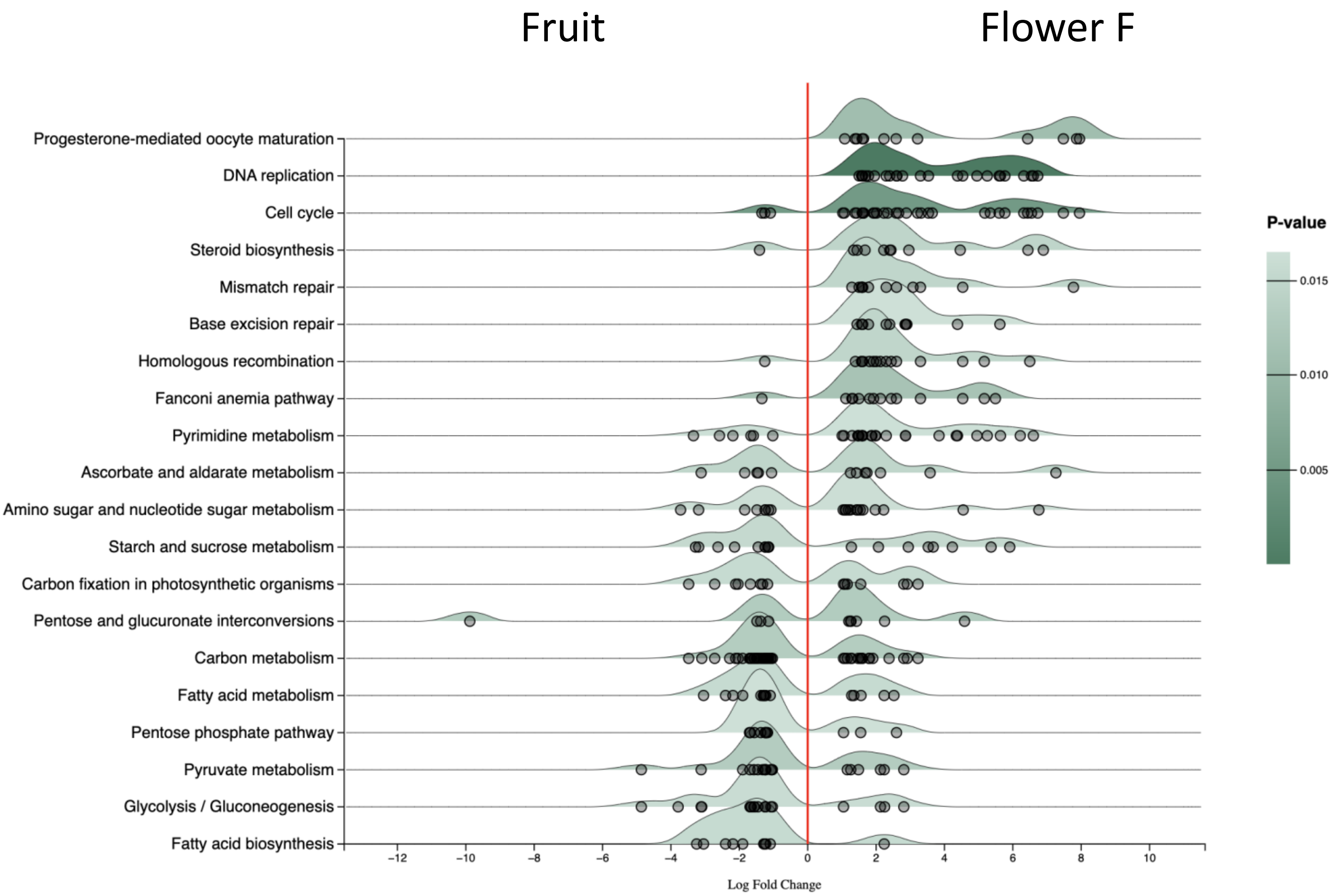


**Supplementary Figure 4.** Ridge plot of KEGG pathways defined by the Over-representation analysis of the transcriptome of fruit and female flower. The circles represent genes in each pathways that differentially expressed between the organs. Pathways are shown in decreasing order of log2 fold change.

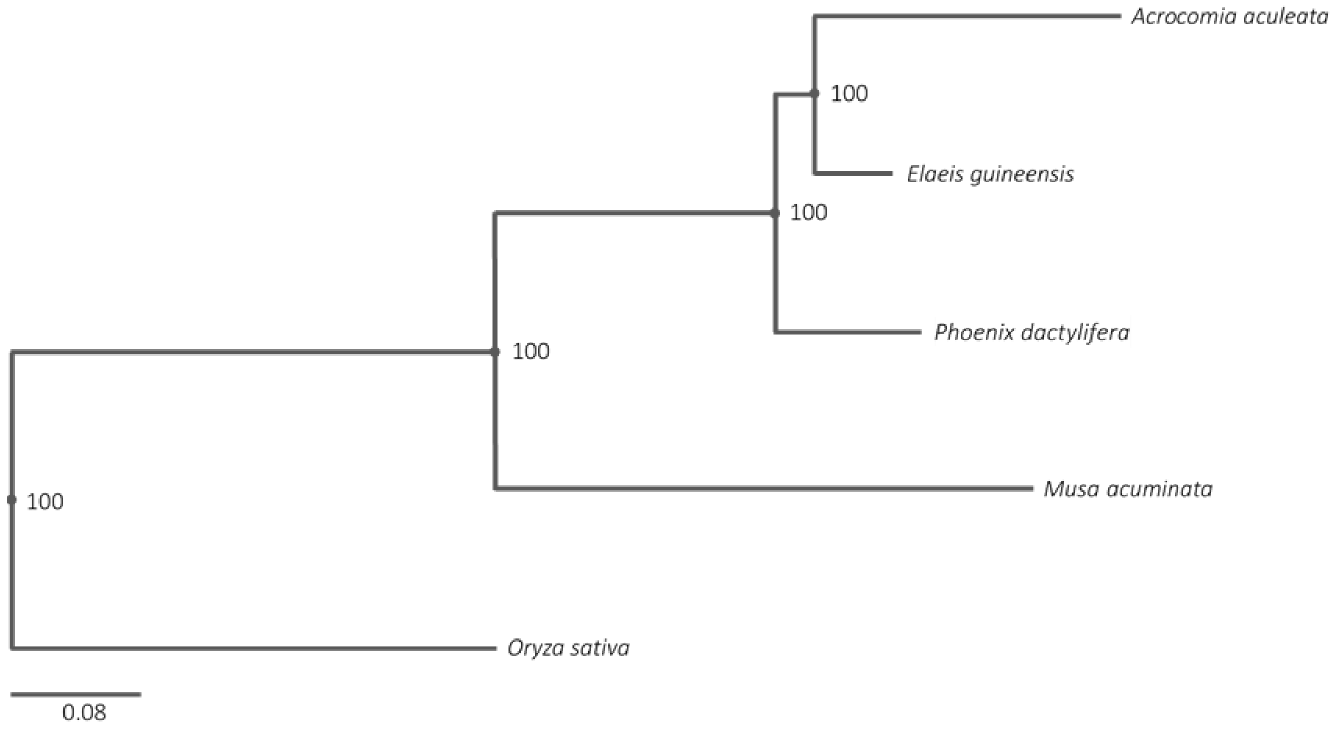


**Supplementary Figure 5.** Maximum likelihood phylogenetic tree based on the concatenated alignment of 1,994 single-copy orthologous proteins conserved in macaúba palm (Acrocomia aculeata), Elaeis guineensis (African oil palm), Musa acuminata (Banana), Phoenix dactylifera (date palm) and Oryza sativa (rice). Numbers at each node are the bootstrap values percentages.
